# Rapid and wash-free anti-drug antibody assay enabled by the complementary nanoluciferase biosensor

**DOI:** 10.1101/2025.02.14.638302

**Authors:** Ruoxuan Sun, Mark G. Qian, Xiaobin Zhang

**Author notes:** Corresponding author: Ruoxuan Sun:, Xiaobin Zhang.

## Abstract

Biotherapeutics have demonstrated remarkable therapeutic efficacy across a wide range of diseases; however, their clinical application is often hindered by the development of anti-drug antibodies (ADAs). Traditionally, ADA monitoring has relied on ligand-binding assays (LBAs) such as enzyme-linked immunosorbent assays (ELISA) or electrochemiluminescence assays (ECLIA). Although effective, these methods require multiple washing and incubation steps, extending data generation time to several hours or even days. In contrast, homogeneous immunoassays offer a promising alternative due to their “mix-and-read” feature, which significantly reduces operational complexity, time, and cost compared to conventional LBAs. In this study, we explored the feasibility of rapid ADA detection using a wash-free homogeneous assay based on the split nanoluciferase (SplitNluc) biosensor system. Through case studies involving diverse therapeutic modalities including antibody-drug conjugates (ADC), Fc-fusion protein, Fc-less multidomain biotherapeutics (MDB), and single-domain antibody (SdAb), we demonstrated that the SplitNluc platform is a versatile and efficient tool for robust and rapid ADA monitoring. This approach not only streamlines the detection process but also holds potential for broader applications in biotherapeutic development and clinical monitoring.

## Introduction

The administration of biotherapeutic agents can result in the production of anti-drug antibodies (ADAs) within the patient population. The presence of ADAs can significantly impact the pharmacokinetics and pharmacodynamics of the therapeutic, and in some cases, has been associated with severe adverse events (1, 2). Therefore, it is recommended by regulatory guidelines and industry white papers to closely monitor the magnitudes of ADAs in nonclinical and clinical stages by multi-tiered bioassays(3–5). ADAs are generally measured by ligand-binding assays (LBAs) such as enzyme-linked immunoassay (ELISA) or electrochemiluminescence assay (ECLIA). In regards of the assay configuration, bridging format is more acceptable, while direct binding is applicable in some cases. However, these assays involve multiple steps, which may take several hours or even days to complete. As so, it will be beneficial to develop a homogeneous, “mix-and-read” serological assay for ADA detection would significantly streamline the workflow, enabling more rapid and efficient assessment of immunogenicity during drug development.

Nanoluciferase (Nluc) is a 19 kDa engineered luciferase derived from deep-sea shrimp *Oplophorus gracilirostris*(6). Using imidazopyrazinone as the substrate, Nluc generates stable luminescence over 150-fold stronger than firefly or *Renilla* luciferases, not to mention its much lower molecular weight. In the study by Nath et al., an ELISA-like bridging ADA assay was developed utilizing Nluc as the reporter(7). Later, Promega established the complementary version of the Nluc platform and commercialized it to meet various research needs(8). This system is based on splitting Nluc into two fragments: LgBiT (derived from strand β1-9) and SmBiT (derived from strand β10). LgBiT and SmBiT exhibit low natural binding affinity, and enzyme activity is restored only when they are brought into close proximity. This system (also termed NanoLuc Binary Technology, or NanoBiT^®^ in short) is widely used for monitoring the protein-protein interaction (PPI) *in vitro* or in live cells, as well as screening PPI modulators(9, 10). By coupling LgBiT and SmBiT with biological probes such as antibodies or oligonucleotides, the NanoBiT^®^ can also be implemented in both direct and competitive format to study a variety of intracellular or blood-derived analytes (11–16). Some reports described the serological quantitation of therapeutic antibodies, pathogen-specific antibodies, and autoantibodies via NanoBiT^®^ system(17–19). To date, the only documented Nluc complementation-based ADA detection was described by Ni *et al*. based on marketed monoclonal antibody drugs(15).

Using the commercially available reagents, we developed a streamlined SplitNluc homogeneous ADA assay that can be applied to quickly evaluate ADA levels in blood samples within about 30 min (Scheme 1). The signal development is based on the close proximity of LgBiT and SmBiT-conjugated drug molecules upon the formation of bridging immunocomplex. The assay showed good sensitivity and precision, and ideal adaptability to biotherapeutics of various sizes. We also investigated the immunogenicity of SdAb using LgBiT/SmBiT fusion protein generated via bioengineering instead of chemical labeling. In general, SplitNluc represents an innovative, convenient strategy for immunogenicity analysis.

**Scheme 1.**
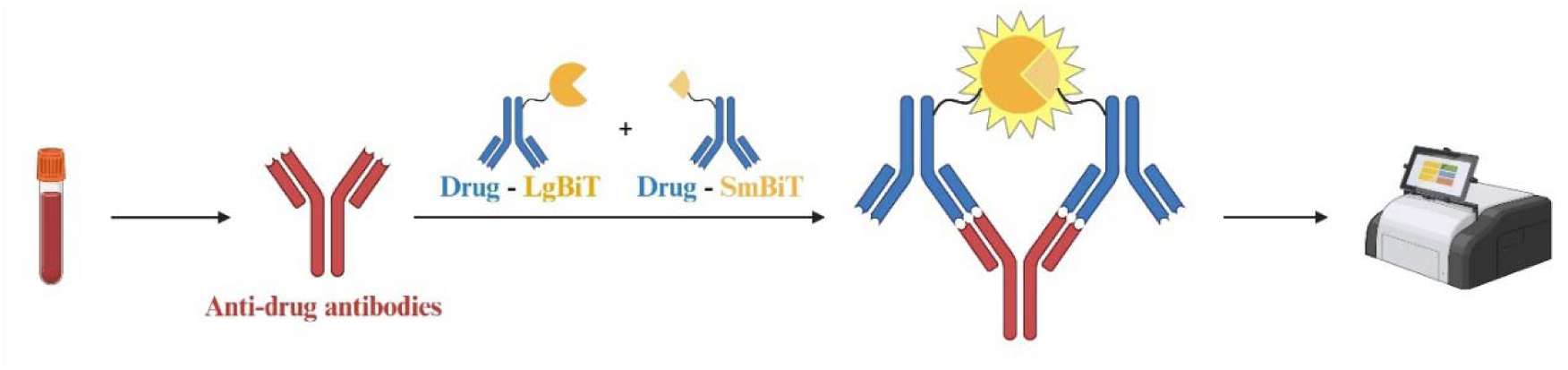
The workflow of SplitNluc assay to detect anti-drug antibodies.

## Materials and Methods

### Materials

The Lumit® labeling kit, substrate A, dilution buffer, LgBiT-labeled anti-mouse antibody were purchased from Promega (Madison, WI, USA). EZ-Link Sulfo-NHS-LC-biotin was purchased from Thermo Scientific (Waltham, MA). MSD GOLD Sulfo Tag NHS-Ester, streptavidin coated MSD plates and 4×MSD buffer T were purchased from Meso Scale Discovery ([MSD] Rockville, MD, USA). Cynomolgus macaque plasma (gender pooled) and human serum (from individual healthy donors) were purchased from BioIVT (Hicksville, NY, USA). Anti-drug antibody 1 (ADC1), immunocytokine 1 (IC1), multi-domain biotherapeutic 1 (MDB1) are proprietary molecules produced internally. Single-domain antibody 1 (SdAb1) and its derivative with *C*-end alanine extension (SdAb1-A) were produced internally based on the sequence 1 from US9573992B2(20). The LgBiT/SmBiT-SdAb1 fusion proteins were provided by Genscript (Nanjing, Jiangsu, China) through cell-free protein synthesis. Polyclonal rabbit antibodies against ADC1 and MDB1, mouse monoclonal antibodies against MDB1 were all produced in house.

### Bioconjugation

The preparation of LgBiT/SmBiT tagged ADC1, IC1 and MDB1 were conducted using Lumit® labeling kit as per the manufacture’s guidance. Successful conjugation was confirmed by native SDS-PAGE gel followed by InstantBlue^®^ (Abcam, Waltham, MA, USA) staining. The conjugated products were stored in 50% glycerol in PBS for storage in 4°C (short term) and 20°C (long term). Biotin- and Sulfo Tag (ST)-conjugated SdAb1 were prepared following the manufacturers’ instructions.

### Animal study

Eight BALB/c mice were intraperitoneally administrated with 10 mg/kg IC1 every four days. The plasma samples were harvested at least 14 days after the first dose. Three mice from vehicle-treated group were also included in this study. The study protocol was approved by the Institutional Animal Care and Use Committee of Takeda Pharmaceutical Co. Ltd.

### SplitNluc assay procedure

5 μl of assay mixture containing LgBiT and SmBiT-labeled drugs (ADC1, IC1, and MDB1) were diluted in Lumit assay buffer and dispensed in low volume 384 well white plates (Greiner, Monroe, NC, USA). Every other column and row are used to prevent signal leakage (totally 96 wells are used in each plate). Subsequently, 5 μl of diluted plasma samples were added to each well and for a 30 min incubation on an orbital shaker set at 25 °C. For signal development, 5 μl of diluted (50-fold) Lumit Reagent A were supplemented and incubate for 5 min to achieve stable signal. For case study 3 involving SdAb, 0.1 μg/ml LgBiT-SdAb1 and 0.05 μg/ml SmBiT-SdAb1 were incubated with equal volume of 100-fold diluted human sera for 30 min. A higher dilution ratio, 1:100, was applied to the substrate to reduce background. Luminescence intensities were acquired on a ID3 multi-mode plate reader (Molecular Devices, San Jose, CA, USA). The reading height and integration time were set to be 6.43 mm and 200 ms, respectively.

### ECLIA procedure

Serum samples were incubated for 1 h with 24-fold diluted 1% BSA/PBST containing 0.1 µg/ml biotin- and ST-tagged IC1 or SdAb1. The mixture was subsequently transferred to a small-spot streptavidin-coated MSD plate pre-blocked with BSA and allowed to incubate for 1 hour to enable formation of the bridging complex. Following multiple washes with PBST, the plates were scanned using a Meso QuickPlex SQ 120MM reader with 150 µl of 2×MSD buffer T. All incubation steps were performed at room temperature with 600 rpm shaking.

### Statistical analysis

All graphing and statistical analysis were performed in Graphpad Prism or Microsoft Excel. The inter-dataset correlation was determined by non-parametric Spearman analysis.

## Results

### Case study 1: SplitNluc ADA assay for an ADC

#### Evaluating the feasibility of SpiltNluc assay for ADA measurement

A previous study has demonstrated the feasibility of Nluc complementation as an ADA detection method(15). However, that assay involves sophisticated bioengineering and photoconjugation, which may not be ideal for widespread use. Upon the recent introduction of the commercialized NanoBiT^®^ antibody labeling kit, we aim to leverage it and develop a refined SplitNluc assay for expediate measurement of ADA. For the initial assessment of the refined SplitNluc assay, we prepared the rabbit polyclonal PC against ADC1 in mouse, cynomolgus monkey, human plasma as well as 0.5% BSA/PBS. LgBiT- and SmBiT-tagged ADC1 were prepared by the lysine-specific, HaloTag-mediated strategy as per the protocol by manufacturer. The successful conjugation of LgBiT-ADC1 and SmBiT-ADC1 was confirmed by the bands with increased molecular weight observed on SDS-PAGE gel (Figure S1). We initiated the pilot study with a simple ’3×5’ protocol by combining 5 μl of assay buffer containing 0.25 μg/ml LgBiT-ADC1 and 0.25 μg/ml SmBiT-ADC1 with 5 μl of undiluted PC sample in a white, low-volume 384-well plate, and incubated for 30 min to allow the bivalent bridging complex to form. Subsequently, 5 μl luminescence substrate was added for signal development. As shown in Figure S2, bioluminescence signal could be generated from multiple matrices at different levels. The luminescent signals increased dramatically for 0.5%BSA/PBS samples, and signal reached a plateau at around 8 μg/ml. However, for plasma samples, the background signals of all matrices were too high and there was barely signal increase with 250 ng/ml PC, indicating that matrix effect is potentially a major hurdle for assay optimization. We then found that a pre-dilution of samples can dramatically reduce the background without altering the signals from high-concentration PC, leading to much better assay sensitivity (Figure S3). A 16-fold dilution of cynomolgus plasma was, therefore, used in the following assay development for case study 1.

#### Optimization of the bridging mix

Due to the absence of washing steps, the sensitivity of complementary Nluc-based immunoassay is largely dependent on the concentration of LgBiT/SmBiT-tagged biosensors. While LgBiT and SmBiT exhibit low binding affinity, high concentrations can still generate background bioluminescence due to random association. Conversely, low probe concentration may lead to unsatisfactory assay sensitivity due to the insufficient formation of bridging complex. Therefore, it is essential to identify the optimal probe concentration to achieve desired assay performance. A pilot study was executed to examine all combinations of 250, 125, 62.5, 31.25 and 15.625 ng/ml LgBiT-ADC1 and SmBiT-ADC1 against NC and 500 ng/ml PC. It appeared in Figure 1A that 125 ng/ml LgBiT-ADC1 and SmBiT-ADC1 yielded optimal assay signal increase (S/N = 2.72). Under this condition, 125 ng/ml polyclonal PC yield an S/N ratio of approximately 1.3 (Figure 1B). Hook effect, which is common in homogeneous immunoassays, was not observed with PC concentrations as high as 16 μg/ml. With the addition of 1.25 μg/ml unlabeled ADC1 in the assay mix, the signal was dramatically inhibited which suggested the specificity of the ADA assay.

**Figure 1.**
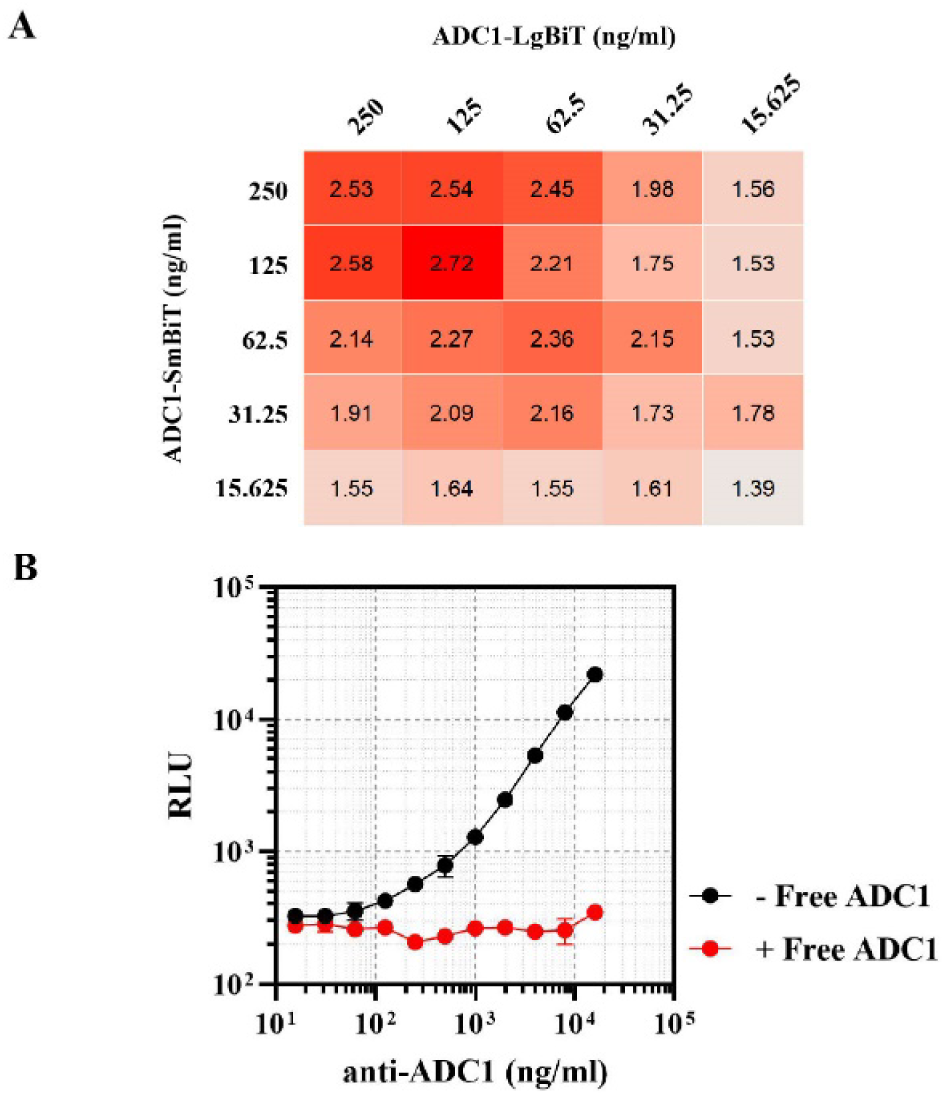
Optimization of SpitNluc assay mix concentration for ADC1. (A) Different concentrations of ADC1-LgBiT and ADC1-SmBiT were combined and used to examine diluted blank pooled cynomolgus plasma (NC) and blank plasma spiked with 500 ng/ml rabbit polyclonal anti-ADC1 (PC). The PC/NC signal ratios are demonstrated in the table. Among all, 125 ng/ml LgBiT-ADC1with 125 ng/ml SmBiT-ADC1 showed optimal result. The heatmap was generated by RStudio. (B) SplitNluc assay for 0-6 μg/ml anti-ADC1 under optimized condition. For the spiked group (red), 1.25 μg/ml unlabeled ADC1 was included in the SplitNluc mix. Mean values ± SD are shown (n=2).

#### Impact of immune reaction and substrate incubation time

The repeated plate washing in conventional LBAs reduces assay background and false positives by nonspecific binding; however, long incubation time is required to enable the formation of stable immunocomplexes that are resilient to washing. Due to the wash-free nature, we assume that a much shorter incubation can achieve desired assay signal in SplitNluc ADA assay. To determine the sufficient assay incubation time for optimal ADA detection, we tested incubation times of 15 min, 30 min, and 60 min. As demonstrated in Figure S4A, incubation for 15 min at room temperature already generates substantial bioluminescence signals, and extended incubation further improves assay performance. The substrate incubation time also contributes to the assay signal as indicated by the dose response curves in Figure S4B. While all curves are parallel, extended incubation time indeed cause signal increases. For luminometers with slower scan rates, precise control of substrate incubation time may help mitigate the intra-well and intra-plate variations.

#### Assay precision

Precision, especially intra-run precision, is a fundamental parameter defining the reliability of bioanalytical assays. In the validation of regulated ADA assays, an intra-assay precision below 20% is usually required from multiple independent runs at several PC levels(3). To evaluate the precision of SplitNluc ADA assay, we repeatedly assayed the anti-ADC1 standard curve once per day for a total of 6 days. According to the data shown in Table 1, the coefficient of variation (CV) values for all concentration points did not exceed 15%. This high level of consistency not only meets regulatory standards but also supports the feasibility of performing the SplitNluc ADA assay in a singlicate manner, thereby reducing reagent usage and operational complexity while maintaining robust and reliable results.

**Tabel 1.** The analysis of data precision of SplitNluc assay. The PC standards were repeatedly assayed once/day for totally 6 days. CV% was calculated based on the mean and standard deviation (SD) of the 6 replicates.

| PC (ng/ml) | Rep 1 | Rep 2 | Rep 3 | Rep 4 | Rep 5 | Rep 6 | Mean | SD | CV |
| --- | --- | --- | --- | --- | --- | --- | --- | --- | --- |
| 0 | 277 | 295 | 302 | 326 | 292 | 292 | 297 | 15 | 5.0% |
| 15.625 | 281 | 319 | 308 | 331 | 297 | 326 | 310 | 17 | 5.6% |
| 62.5 | 315 | 373 | 333 | 389 | 292 | 379 | 347 | 36 | 10.3% |
| 250 | 569 | 696 | 566 | 571 | 488 | 585 | 579 | 61 | 10.5% |
| 1000 | 1414 | 1751 | 1266 | 1481 | 1344 | 1585 | 1474 | 160 | 10.8% |
| 4000 | 5849 | 7543 | 5730 | 6192 | 6536 | 7695 | 6591 | 772 | 11.7% |
| 16000 | 30407 | 30825 | 25956 | 31349 | 30660 | 35638 | 30806 | 2809 | 9.1% |

### Case study 2: SplitNluc ADA assay for a mouse IgG2a-based IC

Fc-fusion proteins such as ICs represents a novel category of bifunctional biotherapeutics that exhibits superior pharmacological value in various conditions especially cancer(21). Herein, we evaluated the SplitNluc ADA assay for IC1 which consists of a mouse IgG2a scaffold and a cytokine genetically fused on the *C*-terminus of the heavy chains. The probe preparation was prepared and confirmed by the same procedure as ADC1 (Figure S5). ADA level was measured for totally 11 samples (three from vehicle group and eight from IC1 group) by both bridging ECLIA and SplitNluc assays. As demonstrated in Figure 2A, both methods successfully distinguished the ADA-positive samples from the negative controls. The results from the SplitNluc assay closely aligned with those from the ECLIA, as evidenced by the strong correlation presented in Figure 2B. The consistency highlights the reliability of the SplitNluc assay as a viable, alternative strategy for ADA detection.

**Figure 2.**
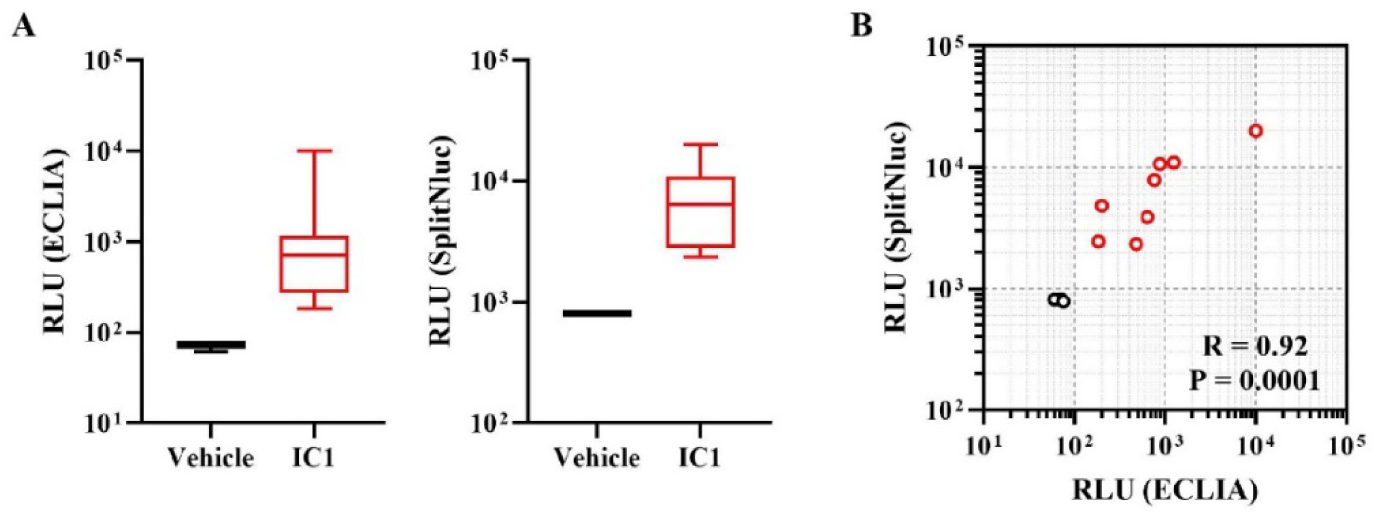
ADA assay results of IC1-treated mice. (A) ECLIA and SplitNluc results of vehicle-(n=3) and IC1-(n=8) treated mice. (B) Correlation of RLU results obtained from two methods. Negative and positive samples are outlined in black and red, respectively. R and P values were calculated by non-parametric Spearman analysis.

### Case study 3: SplitNluc ADA assay for a non-IgG format MDB

Fc-less MDBs represent a critical class of biotherapeutic agent that is composed of multiple domains with distinct functionalities. The method described by Ni et al. involves protein G-mediated photocrosslinking(15), which is not applicable for Fc-less biologics. In this case study, we aimed to interrogate whether ADAs against MDB1 (∼90 kDa), an internal drug candidate, can be measured by SplitNluc platform. The bioconjugation was performed following the standard procedure and confirmed by gel staining (Figure S5). Three PCs (one polyclonal and two anti-idiotype monoclonal) were included here for assay evaluation. As illustrated in Figure 3, all three anti-MDB1 antibodies can be detected through a procedure similar to ADC1, and the addition of free MDB1 resulted in luminescent signal inhibition for all tested PCs. In particular, both mAbs could be detected at higher sensitivities than the pAb. Comparing mAb1 and mAb2, although both PCs yielded ∼ 2.0 S/N at 62.5 ng/ml concentration, the maximal signal amplification of mAb2 was much lower, and it exhibited hook effect at around 4 μg/ml. Prior studies have demonstrated that the sensitivity of anti-drug antibody (ADA) assays is affected by the affinity of the positive control (PC), its epitope profile, and the assay strategy(22), therefore, it is crucial to choose appropriate PCs for ADA assay characterization. Overall, The SplitNluc ADA assay in our study (case 1 and 2) demonstrated the maximal S/N ranging from 10-100 which outperformed the previous study(15). In short, SplitNluc ADA can be leveraged to detect ADAs against non-IgG format modalities.

**Figure 3.**
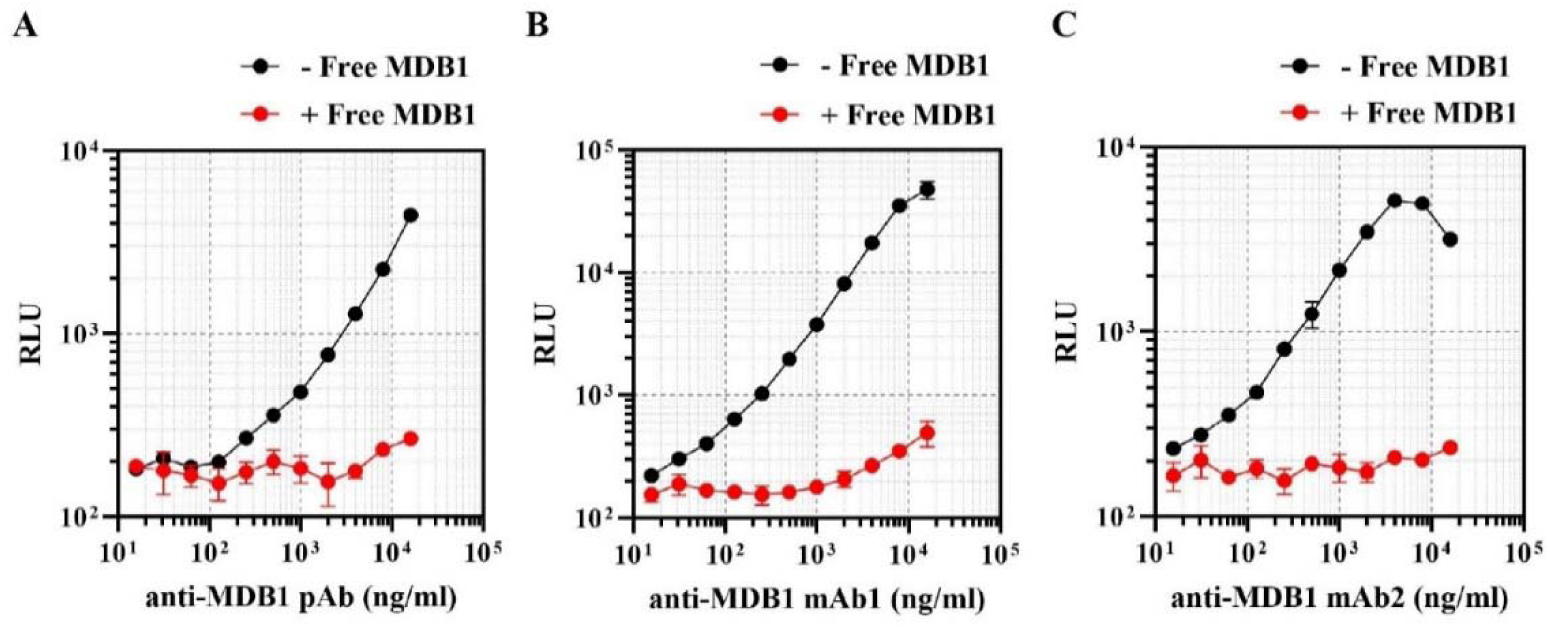
Adaptability of bridging SplitNluc ADA assay to MDB1. Cynomolgus plasma spiked with 0-16 μg/ml anti-MDB1 was analyzed by the SplitNlux mix containing 0.25 μg/ml MDB1-LgBiT and 0.25 μg/ml MDB1-SmBiT. Polyclonal (A) and two monoclonal (B, C) PC were included for assay characterization. For the MDB1 spiked group (dashed lines), 5 μg/ml unlabeled MDB1 was included in the SplitNluc mix before added to samples. Mean values ± SD are shown (n=3).

We also tested another configuration based on the immunocomplex formed by drug + ADA + anti-specie secondary antibody (Figure S6A). To better recapitulate the in vivo situation where the highly abundant endogenous immunoglobulins will compete with ADAs for 2^nd^ antibody binding, we tested the hypothesis in mouse plasma using LgBiT-MDB1, SmBiT-anti-mouse IgG, and mouse monoclonal anti-MDB1 #1 (PC). It appeared that the assay sensitivity was far worse compared to the paratope-mediated bridging format, indicated by the S/N about 1.5 from 1,000 ng/ml PC, which yielded about 8-fold signal increase under conventional format (Figure S6B and Figure 3B). Besides, the Fc-based detection requires specie- and isotype-specific PCs for assay characterization, making it a less ideal option for SplitNluc ADA assay.

### Case study 4: Characterization of pre-existing ADAs against a single domain antibody by SplitNluc assay

Single-domain antibody, also known as a Nanobody or variable heavy domain of heavy chain, is a type of camelid antibody fragment that consists of a single variable domain. The low molecular weight (∼15 kDa) along with the high target binding affinity make it an ideal module to design MDBs. However, the clinical development of SdAb-based biologics is limited by the pre-existing ADAs (PeAbs) which are abundantly found in human as reported(23, 24). Identification of the predominant epitopes will assist the development of next-generation target-binding proteins(25). Strategies such as modifying *C*-terminal sequence and D-amino acid incorporation have been proposed to reduce the immunogenicity of SdAbs(24, 26). Here, we set out to screen PeAbs against SdAb in human sera by both SplitNluc platform and traditional bridging ECLIA, and assess the correlation of two methods. A llama-derived SdAb targeting human serum albumin (designated as SdAb1) was selected for the study. The preparation of SpliNluc probes was not successful using the commercialized labeling kit, resulting in a large amount of free SmBiT and LgBiT which led to high assay background (data not shown). Alternatively, we generated LgBiT- and SmBiT-tagged SdAb through recombinant, cell-free expression platform. It is known that the majority of pre-existing antibodies recognize epitopes located are the *C*-terminal tail of SdAbs(27). As such, the LgBiT and SmBiT were place on the *N*-terminus of the SdAb with a flexible 4×G_4_S spacer (Figure 4A and B). A total of 96 lots of naïve human plasma (50% male, 50% female) were assayed by both SplitNluc assay and ECLIA. The readouts from two methods correlate well (P < 0.0001; R = 0.75, 95% confidence interval = 0.64-0.83) especially for subjects identified as ADA-positive from ECLIA (Figure 4B). As expected, for almost all subjects presenting SdAb1-specific PeAbs, the SplitNluc signals were abolished upon the addition of SdAb1 but not SdAb1-A which contains an additional alanine residue on *C*-terminus (Figure 4D). This result aligns with the previous finding that the majority of PeAbs against SdAb recognizes *C*-terminal epitopes(24). Intriguingly, it appeared that a significantly higher number of subjects which exhibited RLU values under 100 by ECLIA demonstrated high signal intensity in SplitNluc assay (1,000–10,000) (Figure 4C). For instance, subject #55 exhibited only moderate signal magnitude on ECLIA, yet the SplitNluc assay resulted in extremely high signal instead (5^th^ highest in the cohort). This observation indicated potentially the underestimation of PeAb incidence by bridging ECLIA. We hypothesized that due to the absence of washing steps, the low-affinity PeAbs might be better preserved in SplitNluc assay. Nonetheless, more studies need to be conducted to support this hypothesis.

**Figure 4.**
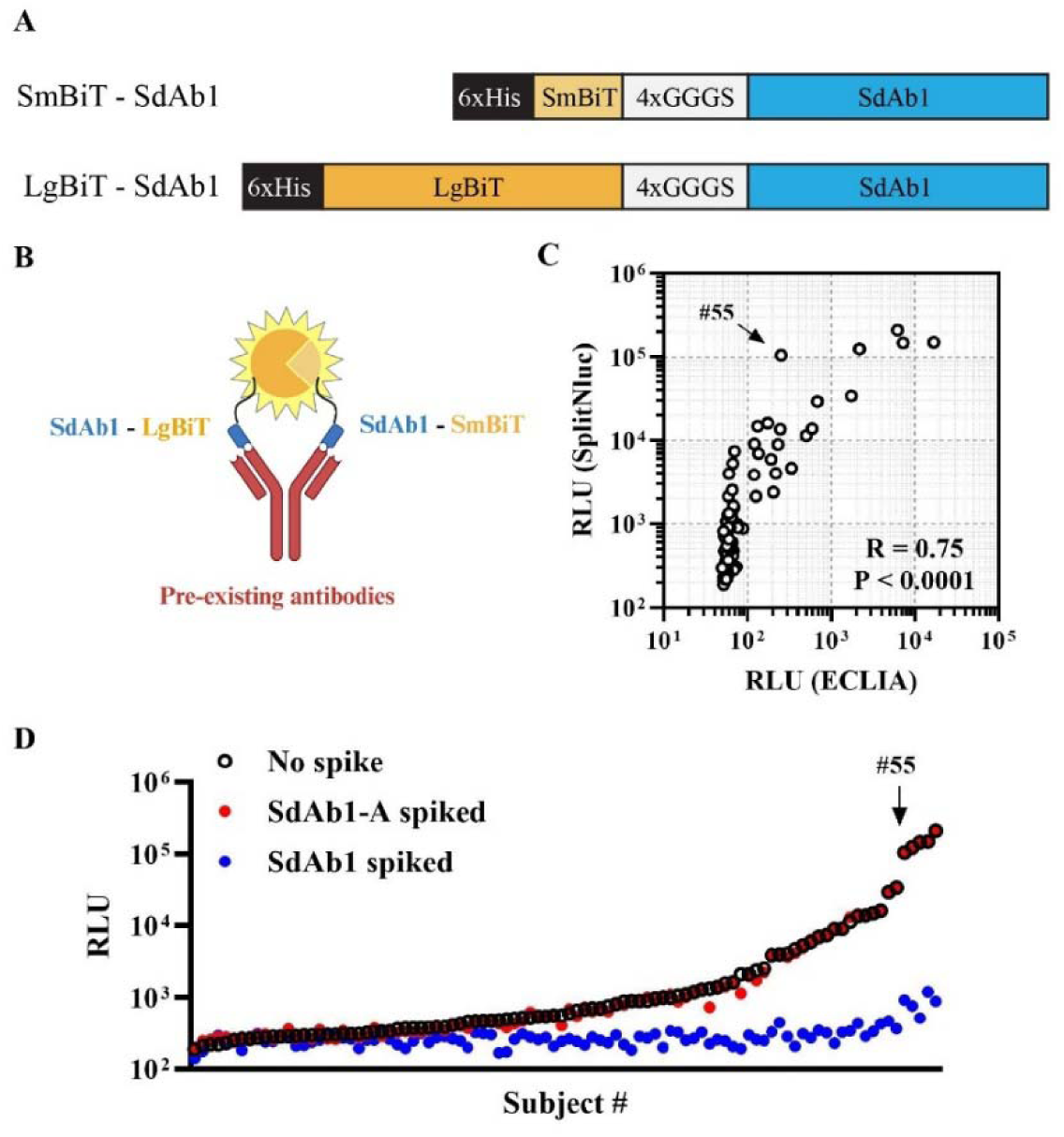
Characterization of PeAb against SdAb in human sera. (A) molecu-lar structure of SmBiT- and LgBiT-tagged SdAb1 used in Split-Nluc assay. (B) Mechanism of PeAb screening by SplitNluc strate-gy. (C) Results of PeAb against SdAb1 from ECLIA and SplitNluc assays. The dot plot showed the correlation between two assays. R and P values were calculated by non-parametric Spearman analysis. (D) SplitNluc assay result of each individual subject with pre-spiked SdAb1 (blue dots) or SdAb1-A (red dots).

## Discussion

The development of safe and efficacious biotherapeutics relies on robust immunogenicity testing and risk mitigation strategies(28). Today, the landscape of ADA assay methodology, especially for regulatory studies, is dominated by heterogeneous assays represented by ELISA and ECLIA. Other methods such as surface plasmon resonance and Gyrolab microfluidic system are also applied but with much lower prevalence (29, 30). These assays either rely on specialized equipment or undergo tedious procedures. To date, amplified luminescent proximity homogeneous assay (AlphaLISA^TM^) technology remains the only wash-free immunoassay accepted by industry(31). Unlike SplitNluc which is based on complementary Nluc system, Alpha technology involves the singlet oxygen-mediated chemiluminescence reaction which is proximity-dependent. However, the protocol for the AlphaLISA-based ADA assay can be somewhat intricate and time-consuming. For the first time, we reported here the convenient wash-free ADA assay based on analyte-triggered complementation of Nluc biosensor. Using ADC1 (case 1) as an example, we described the optimization of SplitNluc assay by modifying sample dilution and probe concentration. We also discussed the impact of immune reaction and substrate incubation time on the assay result. SplitNluc ADA assay also proved itself as a feasible strategy for non-IgG modalities (case 2 and 3). In terms of probe generation, both chemical labeling and recombinant expression can be considered as per the specific scenario. In addition, using SplitNluc as an innovative research tool, we confirmed the previously reported characteristics of PeAb against SdAb in human (case 4). In addition to the ADA detection practices described in this study, the SplitNluc strategy can also be utilized for neutralizing antibody screening and ADA epitope characterization in the wash-free manner.

Despite the advantages such as efficiency and accuracy, some of the drawbacks of SplitNluc ADA assay should also be noted. First, SplitNluc assay has relatively lower sensitivity compared to bridging ECLIA. Besides, the dynamic range of SplitNluc may not be comparable to ECLIA either. These issues could be improved to some level through further tuning, but the limitation is largely attributed to the intrinsically distinct detection efficiency of ECL and bioluminescence. Note that the readout of RLUs varies among different plate readers, meaning that the optimization of SplitNluc-based immunoassays is specific to the given instrument, and parameters such as reading heights and integration time should be carefully considered to achieve optimal results. The system itself may need further optimization on factors such as linker length and labeling strategy to adapt more complicated situations. To overcome the signal variation caused by inconsistent substrate incubation time between wells or plates, the Nluc-fluorescence acceptor fusion protein introduced by Ni *et al.* can be included as a calibrator to convert the absolute RLU to a more accurate ratiometric readout(15). This strategy may further enable the high-throughput, automated assay which may take much more time. Other key considerations such as drug-tolerance and matrix selectivity should also be investigated in the following studies for comprehensive evaluation of this platform. In general, although SplitNluc assay holds promises for its attractive features, it is not anticipated to be broadly applied in regulated studies soon given the aforementioned reasons. Nonetheless, with careful evaluation and optimization of engineered luciferase, substrates, and instrumentation, the SplitNluc platform has the potential to be refined and serve as a viable alternative for ADA detection.

The complementary Nluc system is undergoing rapid advancement. On the foundation of binary Nluc system, Dixon *et al.* established the ternary Nluc system by further splitting β1-9 towards β1-and β9(32). Later, Kincaid et al. developed a ready-to-use NHS-activated β9 and β10 which enabled simple one-step chemical conjugation with detecting antibodies, and paired with genetically evolved β1-8 (LgTrip) for signal development. This advanced strategy is not only more adaptable, but bypassed the HT-bases conjugation which introduced additional steric hindrance that could potentially mask some epitopes. As part of our next steps, it will be insightful to compare the ADA detection efficiency between binary and ternary SplitNluc system. Also, other split enzymes based on β-lactamase, horseradish peroxidase, etc., have been reported, and some are being utilized for serological antibody assays(33, 34). Revolutionary *de novo* enzyme design informed by deep learning has also made progress in the development of luciferase(35). With continuous efforts in enzyme engineering and assay optimization, we are optimistic that the wash-free homogeneous assay will become an invaluable tool in bioanalytical applications.

## Conclusion

By harnessing the complementary Nluc system, we introduced the wash-free SplitNluc platform for rapid ADA detection in blood samples. Compared with traditional ECLIA, our SplitNluc can be completed conveniently by a much shorter workflow (Figure 5). The assay exhibited superior precision owing to the highly simplified workflow which minimizes the artifacts during the whole procedure. We demonstrated the adaptability of this SplitNluc ADA assay for both IgG-based and Fc-less drugs. This strategy can also be utilized to assist the epitope identification and deimmunization of biotherapeutics. In conclusion, the SplitNluc assay represents possibly the best-streamlined platform for ADA detection to date.

**Figure 5.**
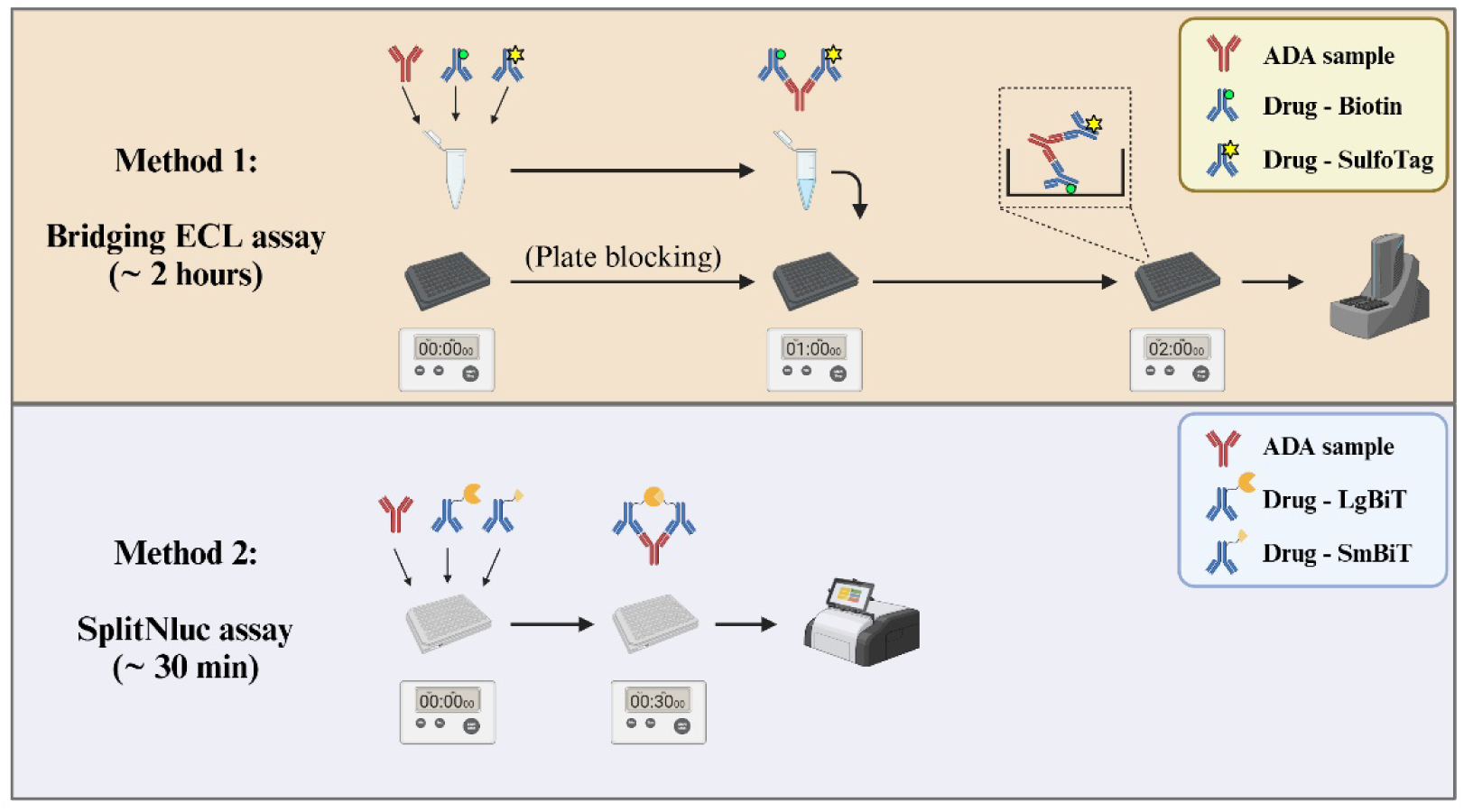
Comparison of conventional bridging ECLIA and wash-free SplitNluc. Method 1: Conventional ADA assay of bridging format detected by MSD system. Method 2: Homogeneous SplitNluc ADA assay based on Nluc complementation.

## Acknowledgement

The authors would like to thank Dongmei Zhang from Global Oncology drug Discovery Unit for providing the animal study samples related to IC1. We also appreciate the support by Dongdong Wang from Global Biologics for providing SdAb1 and SdAb1-A proteins.

## Author Contributions

RS and XZ designed the study. RS acquired and analyzed the data. RS, MGQ and XZ wrote the manuscript. All authors are employed by Takeda Development Center Americas, Inc. The authors declare no further financial conflict associated with this study.

## Supplementary materials

**Figure S1.**
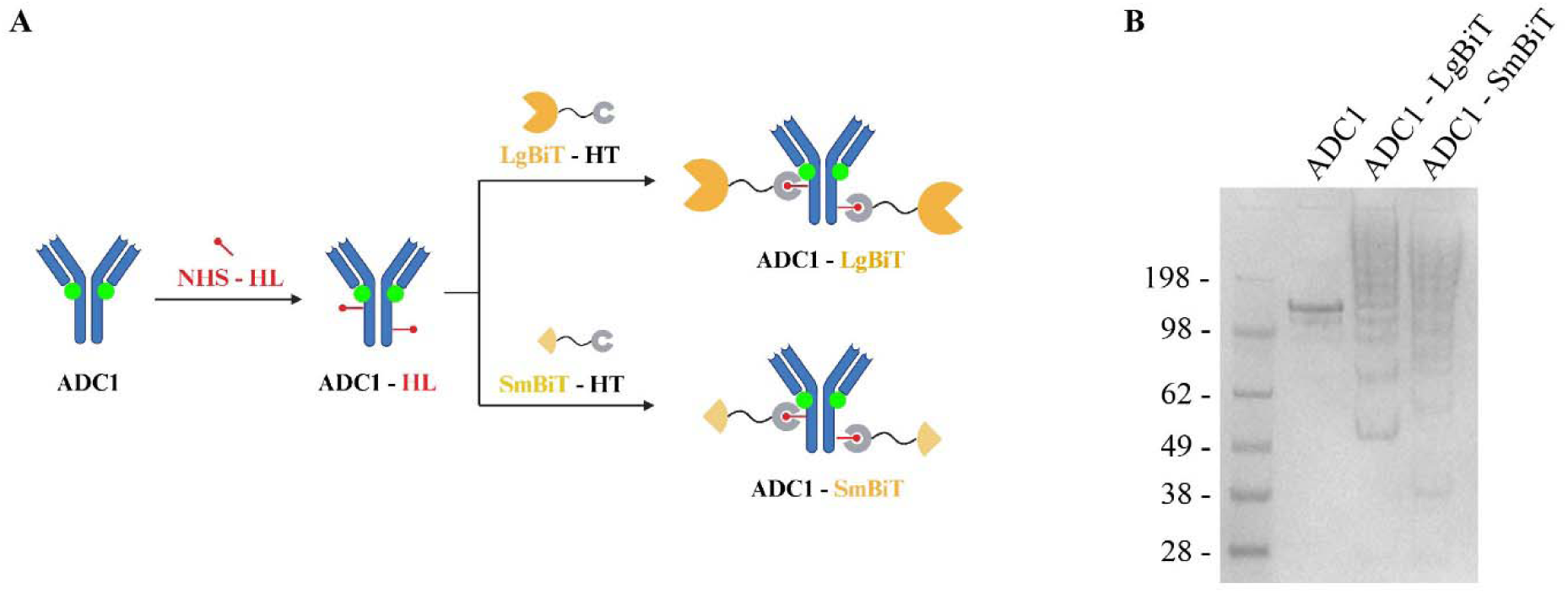
Preparation of LgBiT/SmBiT-conjugated ADC1. (A) The procedure to generate ADC1 with LgBiT or SmBiT tagging. Halo ligand (HL) was first introduced to ADC1 via amine coupling reaction. Subsequently, after removal of free NHS-HL, LgBiT/SmBiT-Halo Tag (HT) fusion proteins were covalently conjugated to HL moieties on ADC1. (B) The validation of ADC1-LgBiT/SmBiT by gel staining (non-reduced). The appearance of high molecular smear represents the formation of conjugated products.

**Figure S2.**
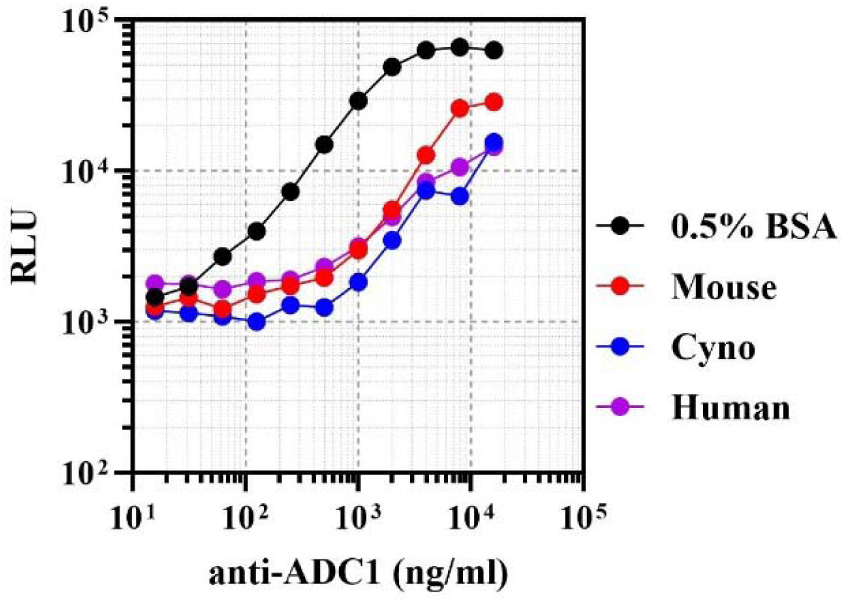
Performance of SplitNluc ADA assay with different matrices. Polyclonal anti-ADC1 (0-16 μg/ml) spiked in 0.5 % BSA/PBS, cynomolgus monkey, human and mouse plasma without dilution were assayed by LgBiT-ADC1 and SmBiT-ADC1.

**Figure S3.**
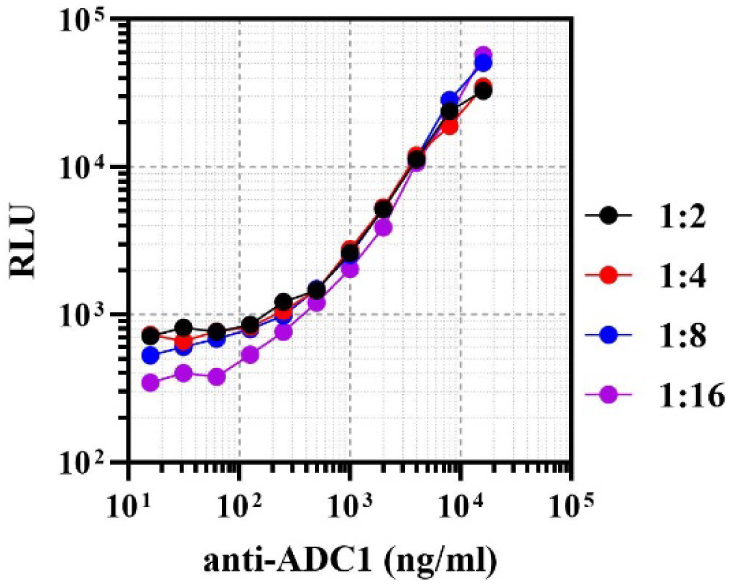
Optimization of sample dilution factor for ADC1 ADA assay. Polyclonal anti-ADC1 (0-16 μg/ml) spiked in cynomolgus plasma was diluted 2, 4, 8 and 16-fold in Lumit assay buffer before analysis by LgBiT-ADC1 and SmBiT-ADC1.

**Figure S4.**
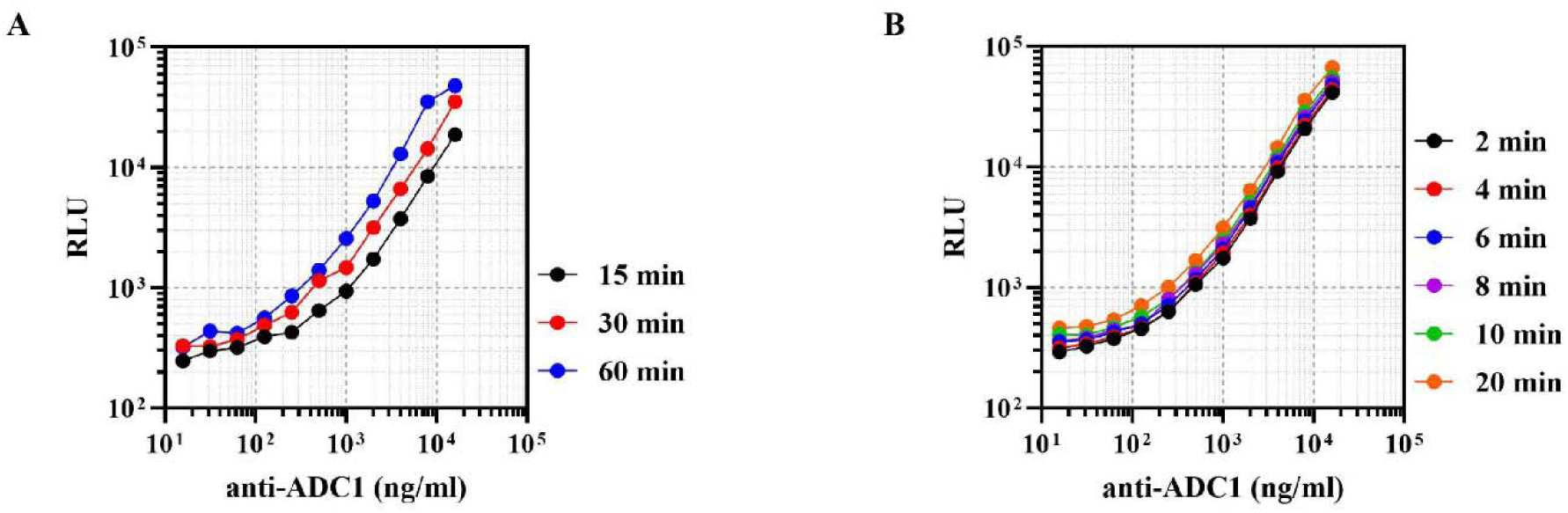
Impact of immune reaction and substrate incubation time to signal intensity. (A) Samples and SplitNluc mix were incubated for 15, 30 and 60 min and the assay performance were compared. 30 min is sufficient to achieve desired signal intensity, while longer incubation further increases the assay sensitivity. (B) Different luminescence substrate time (2–20 min) were compared. Despite the slight increase of luminescence intensities over time, the signal-to-noise remains stable as indicated by the parallel slop of each curve, suggesting the stability of the assay system.

**Figure S5.**
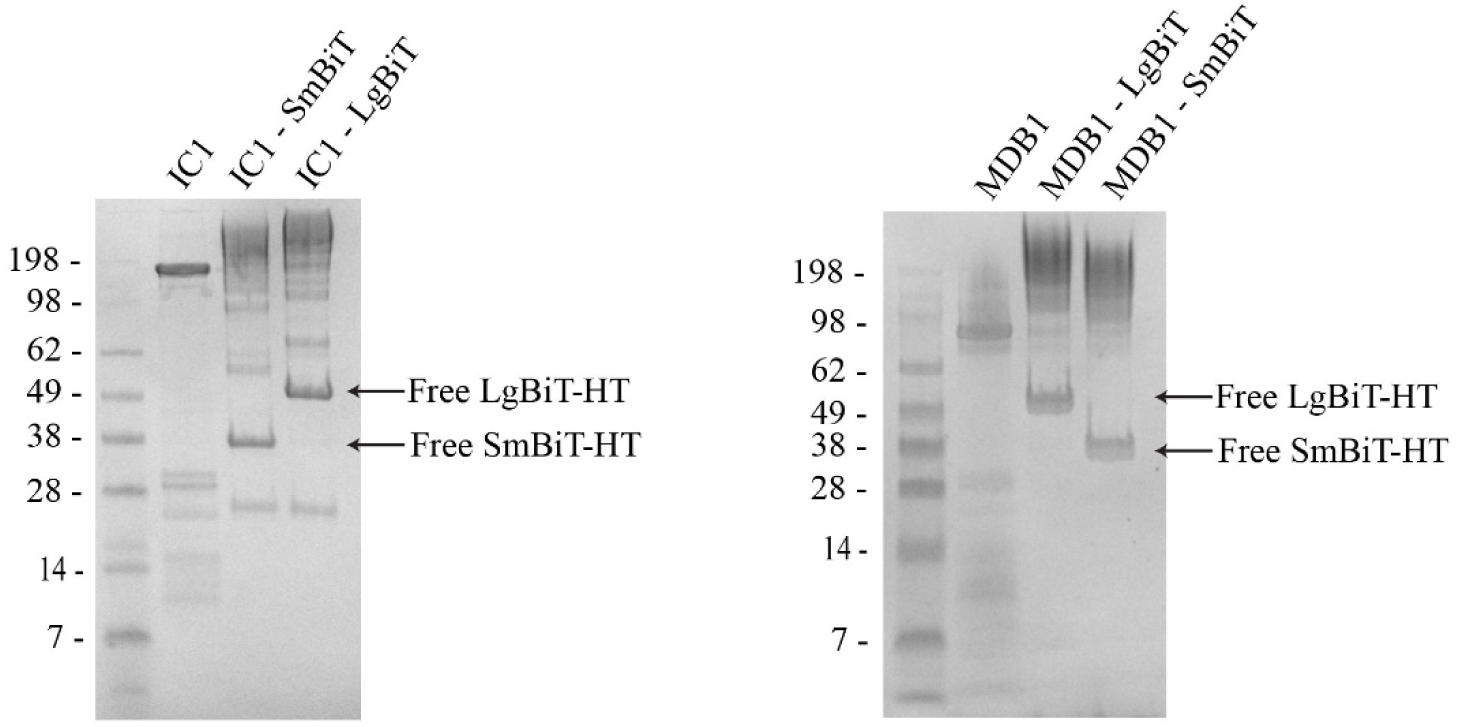
Confirmation of conjugated IC1 and MDB1. Labeled and unlabeled IC1 (left) and MDB1 (right) were analyzed by native gel running followed by staining. Free LgBiT-HT and SmBiT-HT were marked.

**Figure S6.**
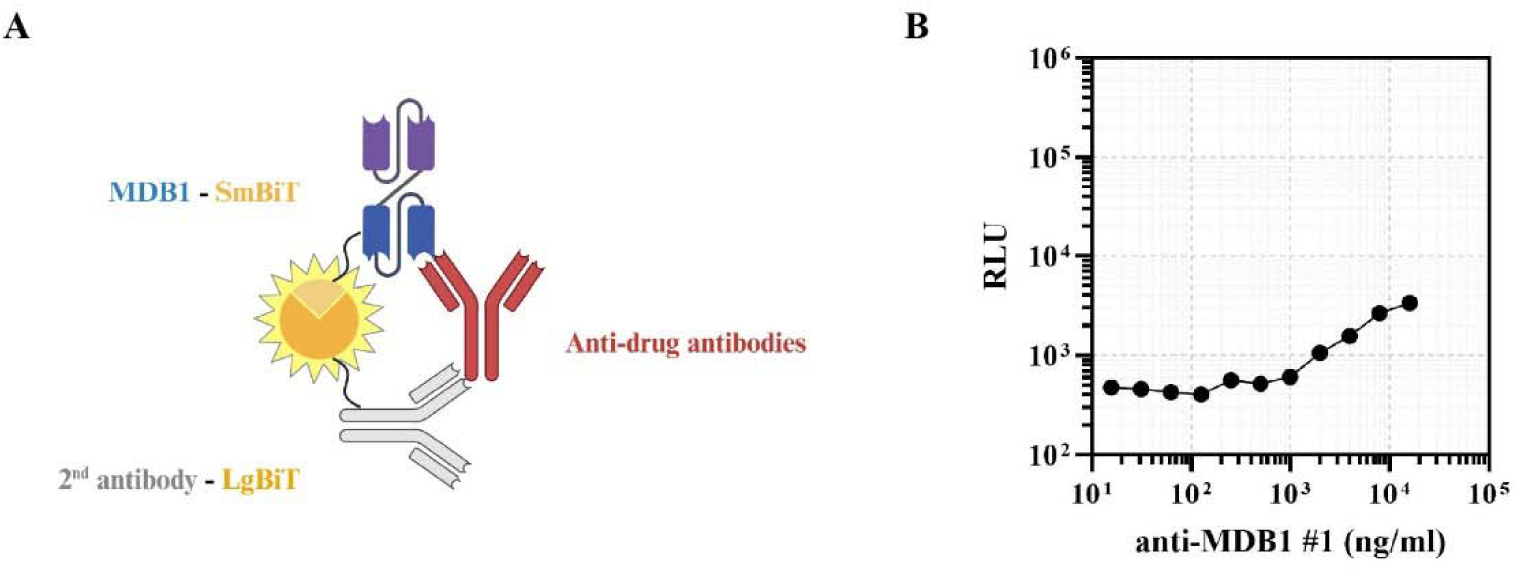
Evaluating an alternative format of SpliNluc ADA assay. (A) The mechanism of luminescent signal generation through the alternative assay format. Note that the real structure of MDB1 is not disclosed and a bispecific T cell engager is used here as an example. (B) Signal intensity of anti-MDB1 #1 under the alternative format.

**Amino acid sequences of LgBiT/SmBiT-SdAb1:**

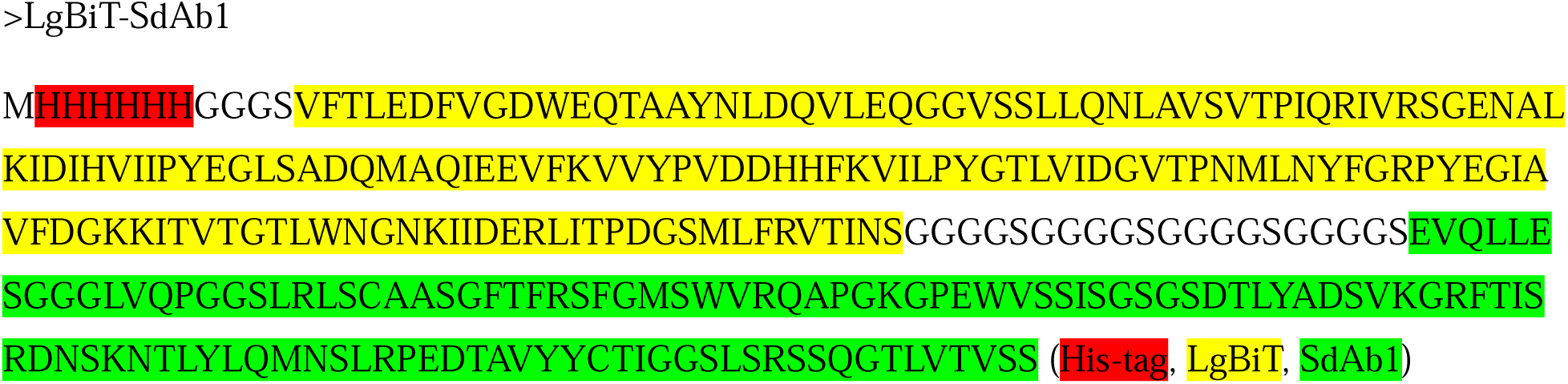
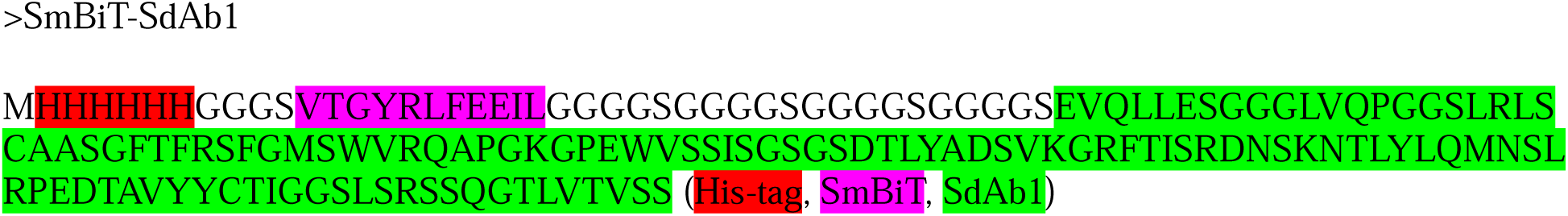

